# Inoculation with a microbe isolated from the Negev Desert enhances corn growth

**DOI:** 10.1101/855528

**Authors:** Noor Khan, Pilar Martínez-Hidalgo, Ethan A. Humm, Maskit Maymon, Drora Kaplan, Ann M. Hirsch

## Abstract

Corn (*Zea mays* L.) is not only an important food source, but also has numerous uses, including for biofuels, fillers for cosmetics, glues, and so on. The amount of corn grown in the U.S. has significantly increased since the 1960s and with it, the demand for synthetic fertilizers and pesticides/fungicides to enhance its production. However, the downside of the continuous use of these products, especially N and P fertilizers, has been an increase in N_2_O emissions and other greenhouse gases into the atmosphere as well as run-off into waterways that fuel pollution and algal blooms. These approaches to agriculture, especially if exacerbated by climate change, will result in decreased soil health as well as human health. We searched for microbes from arid, native environments that are not being used for agriculture because we reasoned that indigenous microbes from such soils could promote plant growth and help restore degraded soils. Employing cultivation-dependent methods to isolate bacteria from the Negev Desert in Israel, we tested the effects of several microbial isolates on corn in both greenhouse and small field studies. One strain, *Dietzia cinnamea* 55, originally identified as *Planomicrobium chinensis*, significantly enhanced corn growth over the uninoculated control in both greenhouse and outside garden experiments. We sequenced and analyzed the genome of this bacterial species to elucidate some of the mechanisms whereby *D. cinnamea* 55 promoted plant growth. In addition, to ensure the biosafety of this previously unknown plant growth promoting bacterial (PGPB) strain as a potential bioinoculant, we tested the survival and growth of *Caenorhabditis elegans* (a test for virulence) in response to *D. cinnamea* 55. We also looked for genes for potential virulence determinants as well as for growth promotion.

## Introduction

Corn (*Zea mays* L.) belongs to the grass family Gramineae, and is an important food, feed, and fuel crop across the globe, especially in the US Midwest (Butler et al., 2018). Corn is both highly productive and strongly influenced by temperature (Hu and Buyanovsky, 2003; Schlenker and Roberts, 2009; Lobell et al., 2013; Prasad et al., 2018) and salinity (Farooq et al., 2015; Iqbal et al., 2018). Therefore, it is critical to develop efficient strategies that improve stress tolerance in this crop and also overcome the significant yield losses predicted to occur under changing soil and climatic conditions. One sustainable solution is to employ plant growth-promoting bacteria (PGPB), which colonize plant roots and improve plant health and growth either directly or indirectly (Backer et al., 2018). Various studies focusing on the beneficial role of PGPB on different crop plants have been published (Vejan et al., 2016; Timmusk et al., 2017; and references therein). However, not all the potentially useful strains that show promise in the laboratory have succeeded in the commercial market mainly because they often lack certain key characteristics that render them abiotic-stress tolerant so that they can persist under challenging environmental conditions. Hence, there is a pressing need for abiotic stress-tolerant PGPB to alleviate the adverse effects of fluctuating conditions in soil.

The present study is a step in that direction. Here, the effects of abiotic stress-tolerant *D. cinnamea* 55 inoculation on the growth and health of corn were evaluated. To our knowledge, this is the first report of growth enhancement in corn mediated by *Dietzia cinnamea*. This hitherto unrecognized PGPB will be useful for further investigations into replacing synthetic amendments with beneficial bacteria.

## Materials and Methods

### Sample collection, bacterial isolation, and culture conditions

To isolate abiotic stress-tolerant PGPB, we screened various bacteria that were previously isolated from the Negev Desert, Israel. Briefly, samples were collected from various sites and pooled to form a composite sample. This mixture was immediately transported to the laboratory for isolation and identification of cultivatable rhizosphere bacteria. The details of the sampling sites, isolation procedure, and the media used for cultivation of the bacterial strains have been described (Kaplan et al., 2013).

### Screening for abiotic stress tolerance abilities

Preliminary screening included the assessment of bacterial strains for their ability to tolerate abiotic stresses by streaking them on Luria-Bertani (LB) agar plates supplemented with NaCl (2, 4, and 6% w/v) to test for salinity stress; PEG (polyethylene glycol; aver. mol. wt. 3,350) at 30, 45, and 60% for drought stress, and pH 4 and 9 for pH stress. Media plates that contained neither salt nor PEG at pH 7.0 served as the controls. The growth of the bacterial strains at high temperature was also assessed by placing a set of streaked plates in a Fisher Scientific Incubator (Model 655D) adjusted to 37°C. For all the experimental sets, plates incubated at 30°C served as the control and bacterial growth was observed over a period of 10 days.

The abiotic stress tolerance of the selected strain was assessed in shake-flask conditions by growing the individual bacterial cultures under control (0% NaCl, pH 7, 30°C), saline (2, 4 and 6% w/v NaCl, pH 7, 30°C), and drought (0% NaCl, pH 7, 30, 45, and 60% PEG 3350, 30°C) conditions in 150 ml Erlenmeyer flasks containing 50 ml LB, with an initial inoculum of about 10^7^ CFU/ml. The flasks were incubated in a New Brunswick Scientific Co. (Edison, NJ, USA) Series 25 incubator shaker at 180 rpm. Viable cells (CFU/ml) were counted at various time intervals for up to 15 days by serial dilution plating on LB agar plates in triplicate (Mishra and Nautiyal, 2012).

Physiological characterization of the selected strain was performed for various plant growth promotion abilities including the production of cellulase, pectinase, xylanase, and protease following standard protocols (Khan et al., 2018). Siderophore production and phosphate solubilization activity were tested on CAS agar and PVK plates respectively, following the methods described by Schwartz et al. (2013).

### Amplification, Purification, and Sequencing of the 16S rRNA Gene

The identification of the bacterial strains was based on phylogenetic and molecular evolutionary analyses (Adhikari et al., 2015). Bacteria were suspended from a single colony grown on LB agar plates into 20 µL of sterile distilled water (SDW). The gene for 16S rRNA was amplified by PCR using the forward primer fD1 and the reverse primer rD1 (Weisberg et al, 1991). Amplification was performed in a total volume of 25 µL containing 14.9 µL SDW, 1 µL of bacterial lysate sample, 2.5 µL of 10× Taq Buffer (MgCl_2_), 0.5 µL fD1 and rD1 (10 µM), 0.5 µL dNTPs (10 mM), 5 µl Q solution and 0.125 µL Taq DNA polymerase. Amplified 16S rDNA products were visualized with ethidium bromide both in the gel and in the gel electrophoresis running buffer and purified from a 0.8% low-melting point agarose gel (100 V, 400 mA, 1 h). The gel extraction was performed with the Invitrogen Quick Gel Extraction Kit according to the manufacturer’s directions. The samples were then sent to Laragen Inc. (http://www.laragen.com/) for further processing and sequencing. The nucleotide sequences were compared against nucleotide databases using the NCBI BLASTn and EzTaxon server 2.1 programs to identify the closest known taxa.

### Phylogenetic analysis

The 16S rRNA gene along with their closest homology sequences were aligned using multiple sequence alignment program CLUSTAL W algorithm implemented in MEGA 6 software by using default parameters. The phylogenetic tree was constructed by Maximum Likelihood method using the MEGA 6 program and evolutionary distances were computed with the help of Kimura’s 2 parameter models. The bootstrap analysis with 1000 replications using p-distance model was performed based on the original dataset to estimate the confidence of a particular clade (Adhikari et al., 2015)

### Plant growth promotion analysis

#### Greenhouse trials

Greenhouse experiments were conducted in pots with corn plants at the Plant Growth Center, UCLA. Seeds of *Zea mays* L. (Golden Bantam), obtained from Baker Creek Heirloom Seed Company, Mansfield, MO, United States, were surface-sterilized by immersing them in 70% ethanol for 1 min, followed by three rinses with SDW. For the treatments, the seeds were bacterized for 3 h by imbibing them in a bacterial suspension, which had been grown for 48 h to contain approximately 10^9^ CFU/ml. Seeds imbibed in LB medium served as the control. The treatments consisted of control and strain 55-treated plants grown in sterile Sungro Potting Mix containing mainly Canadian sphagnum peat moss along with a small fraction of coarse perlite and dolomitic limestone (http://www.sungro.com/professional-product/sunshine-mix-2/). Eight replicates of each treatment, with four plants in each pot, were prepared. Soil moisture was maintained to approximately 20% with water. Plants in all the treatments were grown in parallel and harvested at the same time after 45 days of sowing. Measurements on morphological parameters, namely shoot length and root length, were recorded at the time of harvesting, using general methods as described earlier (Nautiyal et al., 2006a). Plant dry weight measurements were made after drying the plants in a 60°C oven. Photographs of the plants were taken with a Canon PowerShot ELPH350HS camera. The trial was repeated three times and the data were generated from the pooling of all the trials. The data are presented as mean ± standard deviation (SD). The statistical analysis was performed using GraphPad Prism software version 5.01 (GraphPad Software, San Diego, CA, USA).

#### Field studies

The plant growth stimulatory effect of strain 55 was also examined in an outside garden. As a pilot experiment, a pot trial on the pattern similar to the greenhouse trial was set up and the pots were placed in an open, but protected hoop house for examining whether the corn growth promotion mediated by strain 55 was effective under real-life environmental conditions. On observing a positive response of strain 55 on the growth of corn in the pots, we tested the effectiveness of strain 55 by sowing bacterized and untreated seeds directly in the soil of the experimental microplot (2 m x 2 m) within the confines of the hoop house in an outside garden adjacent to the Plant Growth Center, UCLA, Los Angeles. The seeds were planted in four rows, two for each treatment. Each row contained 12 plants, with an intra- and inter-row spacing of about 20 and 60 cm. Harvesting was carried out after 120 days of sowing, followed by the recording of the plant data. The outside experiment was repeated three times in all including the pilot experiment, and the data presented results from the pooling of all trials. The data are presented as mean ± standard deviation (SD). The statistical analysis was performed using GraphPad Prism software version 5.01 (GraphPad Software, San Diego, CA, USA).

#### Determination of whether *D. cinnamea* 55 is virulent

Because soil bacteria may harbor biochemical pathways with pathogenic potential, strain 55 was monitored for its effects on the model nematode worm *Caenorhabditis elegans* and its genome was also examined for the possible presence of virulence genes (see Martínez-Hidalgo et al., 2018).

#### Nematode assays

The activity of *C. elegans* fed with strain 55 under slow-killing conditions was assayed as described by Vílchez et al. (2016). Briefly, the test bacteria were spread on two nematode growth media (NGM) plates and incubated at 30°C for 24 h. Each plate was seeded with a known number of nematodes from the original control plate (*Escherichia coli* OP50), which was determined using a Zeiss microscope at 10X magnification (Carl Zeiss, Oberkochen, Germany). This number served as a zero-h reading. After counting, the plates were incubated at 25°C and scored for nematode death every 24 h for 5 days. The strain *E. coli* OP50 served as a control for estimating the natural death rate of the nematodes and *Pseudomonas aeruginosa* PA14 was the positive control for pathogenicity. The experiment was conducted three times with two replicates of the bacterial strain. The evaluation of the effect of bacteria on *C. elegans* was conducted based on the pathogenicity score given by Cardona et al. (2005). The authors established that a given strain could be designated pathogenic for the nematode if one of the following criteria were met: (i) a diseased appearance at day 2, which included reduced locomotive capacity and the presence of a distended intestine; (ii) percentage of live nematodes at day 2 ≤ 50%; and (iii) total number of nematodes at day 5 ≤ 50%. The presence of any one, two, or three of these criteria was scored to differentiate mild from severe infections. A pathogenic score (PS 1, 2, or 3) was given based on the number of criteria met. A strain was considered non-pathogenic when no symptoms of disease were observed (pathogenicity score, PS 0). Additionally, the influence of the bacteria on movement and propagation of the nematodes was monitored for 120 h. The data are presented as mean ± standard deviation (SD). The statistical analysis was performed using GraphPad Prism software version 5.01 (GraphPad Software, San Diego, CA, USA).

#### Genome sequencing and bioinformatics information

Strain 55 was sequenced at the JGI-DOE as a part of the CSP 1571 project: “The nodule microbiome from legumes living in diverse environments: What other endophytes live within legume nodules besides the nitrogen-fixing symbiont?” The goal of the project was to determine not only the identities of the endophytic bacteria within legume nodules but also the bacteria closely associated with the plant rhizosphere. Some features of the genome are presented in Table 4 and Fig. 5.

The *Dietzia cinnamea* 55 genome was queried using IMG/MER (https://img.jgi.doe.gov/cgi-bin/er/main.cgi) against other sequenced *Dietzia* species and strains as well as other Gram-positive species including *Micromonospora aurantiaca* ATCC 27029 and L5 (Hirsch et al., 2013), *Bacillus subtilis* 30VD-1 (Khan et al., 2018), and *B. simplex* 30N-5 (Maymon et al., 2015) for plant-growth promoting traits such as phytohormone production, abiotic stress resistance, siderophore synthesis, and salinity tolerance. In addition, we queried the genomes for genes encoding virulence determinants such as the type VII secretion system (T7SS), which is frequently found in Gram-positive microbes, especially pathogenic species.

#### Growth conditions and genomic DNA preparation

Strain 55 cells were grown in 5 ml of LB at 30°C for 18 h at 180 rpm. DNA extractions were performed using Invitrogen’s Purelink TM Genomic DNA Mini Kit. The purified DNA was monitored for integrity by gel electrophoresis, and then sent to the US Department of Energy Joint Genome Institute for sequencing.

## Results

### Strain Isolation

Strain 55 was isolated from rhizosphere soil under the canopy of the dominant plant *Zygophyllum dumosum*. During the collection period, the total monthly rainfall was low except in January 2010, normally the rainy season, at which time the rainfall increased to 81.2 mm, which is higher than normal. The following year, in February 2011, the rainfall level was only 21.2 mm indicating that bacteria and plants living in this location have to adapt to significant environmental stress. In addition, the temperature in the driest months was 45-50°C. Based on these parameters, we reasoned that strategies for microbial survival would reflect a high percentage of genes for stress and osmo-tolerance and hypothesized that bacteria with these traits might serve as potential inoculants under conditions of climate change.

### Identification of abiotic stress-tolerant strain 55

*In vitro* experiments conducted on plates demonstrated that strain 55 is an efficient salt-, pH-, and drought-tolerant bacterial species compared to the 40 other bacterial isolates tested (data not shown). Also, several of the other isolates were found to be potential pathogens after 16s rDNA sequencing. Molecular characterization based on 16S rDNA sequence analysis of strain 55 indicated that its closest phylogenetic relationship is to *Dietzia cinnamea* with 99% homology (Fig. 1). A growth curve of strain 55 in presence of 2, 4, and 6% NaCl was monitored at 30°C to up to 15 days. Strain 55 survived mild salt stress conditions with a final CFU of ∼10^7-8^ CFU/ml on day 15. However, a reduced CFU (∼10^5-6^ CFU/ml) was observed in 6% NaCl. A similar trend was recorded in 30, 45, and 60% PEG over a period of 15 days, where strain 55 exhibited a CFU of 10^5-6^ CFU/ml for all three concentrations of PEG tested (Fig S1). Also, strain 55 was found to be sensitive to higher temperatures because the plates incubated at 37°C demonstrated no growth in comparison to the control plates incubated at 30°C, which showed good growth.

**Fig 1.**
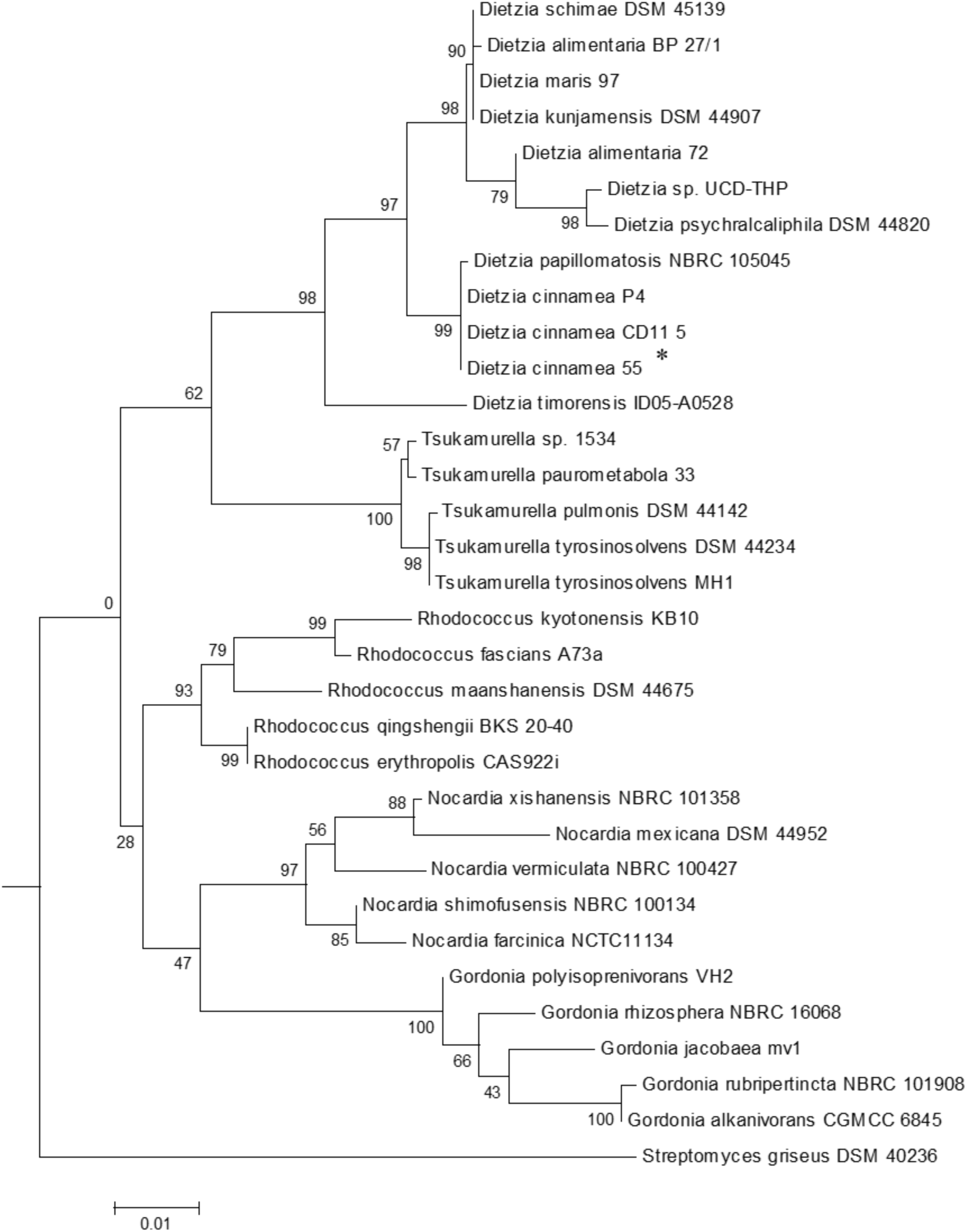
The phylogenetic tree of *Dietzia cinnamea* 55 constructed based on 16SrDNA sequence. Bootstrap values are based on 1000 replications. The asterisk indicates the strain for which the 16S rDNA sequence was determined in this study. Bar, 0.01 substitutions per site.

Among the plant-growth promotion attributes that influence endophytic entry into the plant as well as antagonistic behavior against other microbes, strain 55 tested positive for cellulase, xylanase (Fig. S2), protease, and amylase production, but not for pectin degradation. Biochemical assays for the presence of siderophores and the ability to solubilize phosphate demonstrated that strain 55 performed the former (Fig. S2), but not the latter (data not shown). These conclusions were confirmed by finding genes that code for the production of siderophores in the strain 55 genome (Table 4), but none for phosphate solubilization.

### Adaptations to stress

Maize is well adapted to high temperatures, but grows poorly if severely stressed by drought. Genes with the potential to confer temperature and desiccation tolerance were predicted to reside in the *D. cinnamea* 55 genome. The genomes of two Gram-positive spp. that function as PGPB (*Bacillus subtilis* 30VD-1 and *Micromonospora aurantiaca* ATCC 27029 (https://img.jgi.doe.gov/cgi-bin/er/main.cgi) were used to probe the *D. cinnamea* genome for homologous genes. Possible mechanisms for stress tolerance include the expression of genes for trehalose synthesis and glycine/betaine production, heat shock proteins, and many other functions.

Trehalose, two glucose molecules joined together via an alpha-(1,1) glycosidic linkage, is produced by a wide range of organisms and is important for protecting cells from desiccation (Streeter and Gomez, 2006). Genes for 3 distinct pathways were uncovered in the *D. cinnamea* 55 genome: 1) the MOTS pathway where trehalose synthase (*treXYZ* genes) convert maltooligosyl saccharides are converted to maltooligosyl trehalose; 2) the TPS pathway where in the first step, trehalose-6-phosphate is synthesized from UDP-glucose and glucose-6-phosphate upon the expression of *otsA* (trehalose-6-phoshate synthase) followed by trehalose 6-phosphate phosphatase via *otsB* expression to generate trehalose; and 3) the synthesis of trehalose from maltose directly via *treS* (TS: trehalose synthase). Genes for all three pathways were detected in *D. cinnamea* 55. For the MOTS pathway in the *D. cinnamea* 55 genome, *otsB* was not adjacent to *otsA* as it is for *M. aurantiaca* ATCC 27029 (Fig. 5), which occurs in *Rhizobium* and *Paraburkholderia* spp. (Angus et al., 2013).

Several genes encoding stress-related proteins including catalase and peroxidase, the universal stress protein A (UspA), and superoxide dismutases were also detected in the *D. cinnamea* 55 genome. In addition, genes encoding proteins for an osmoprotectant transport system were found in the strain 55 genome (Table 4) as well as genes important for biofilm formation such as the *tad*/*pil* operon (Pu and Rowe-Magnus, 2018). The biofilm state helps plant-associated bacteria tolerate drought stress (Timmusk et al., 2013).

### Effect of *D. cinnamea* 55 on the growth of corn

In a greenhouse experiment performed for 45 days, the effect of corn seed bacterization with strain 55 was assessed in comparison to an untreated control. Plants raised from the treated seeds showed improved overall health that was observed in terms of a significant increase of ∼30% and 16% in plant parameters, namely shoot and root length as well as a ∼70% increase in dry plant biomass over the untreated controls. Also, more extensive root growth was observed in the strain 55-treated plants in comparison to those of control plants (Fig. 2; Table 1). After 35 days, the treated plants (right) exhibited significantly much more root and shoot growth than the control plants.

**Fig 2.**
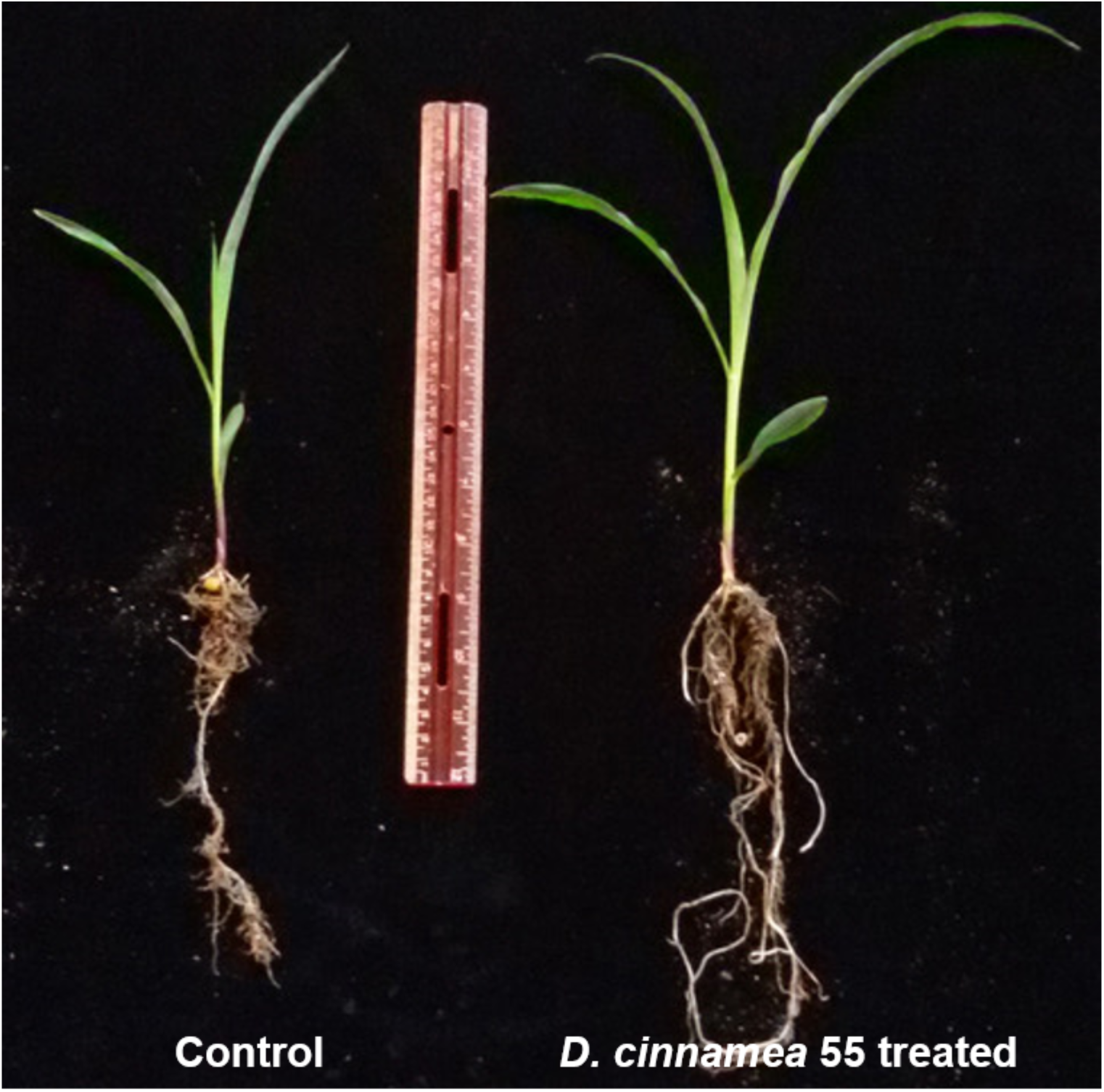
Effect of *D. cinnamea* 55 on the plant growth promotion of corn (Pot trial)

**Table 1.** Effect of *D. cinnamea* 55 inoculation on the growth of corn plants (Pot trial)

| Parameters | Treatments |  |
| --- | --- | --- |
|  | Control | Treated |
| Shoot length (inches) | 32.08±4.62<sup>a</sup> | 41.94±5.54<sup>b</sup> |
| Root length (inches) | 16.63±3.99<sup>c</sup> | 19.34±2.49<sup>d</sup> |
| Dry plant biomass (g) | 10.09±2.93 | 17.70±4.95 |

Values are mean of 8 replicates ± SD of 30 plants in each treatment at the time of harvesting (data recorded after 45 days of experiment setup). Statistical significance of data was validated using One-way ANOVA with Tukey's post hoc test and multiple comparison procedure. Different letters represent values that differ significantly, $P < 0.05$ . Plant biomass data was statistically evaluated using unpaired t-test, $P < 0.05$ .

### Outside garden studies

The effect of inoculating strain 55 on corn plant growth was also assessed in the outside garden experiments. The effectiveness of the bacterial treatment was evident by the stimulation of plant growth parameters where an increase of 39% and 36% was recorded in shoot length and plant dry weight of seeds bacterized with strain 55, respectively, as compared to non-bacterized seeds. Also, a 62% increase in the fresh weight of harvested cobs was observed for treated plants over the control sets (Fig. 3; Table 2).

**Fig 3.**
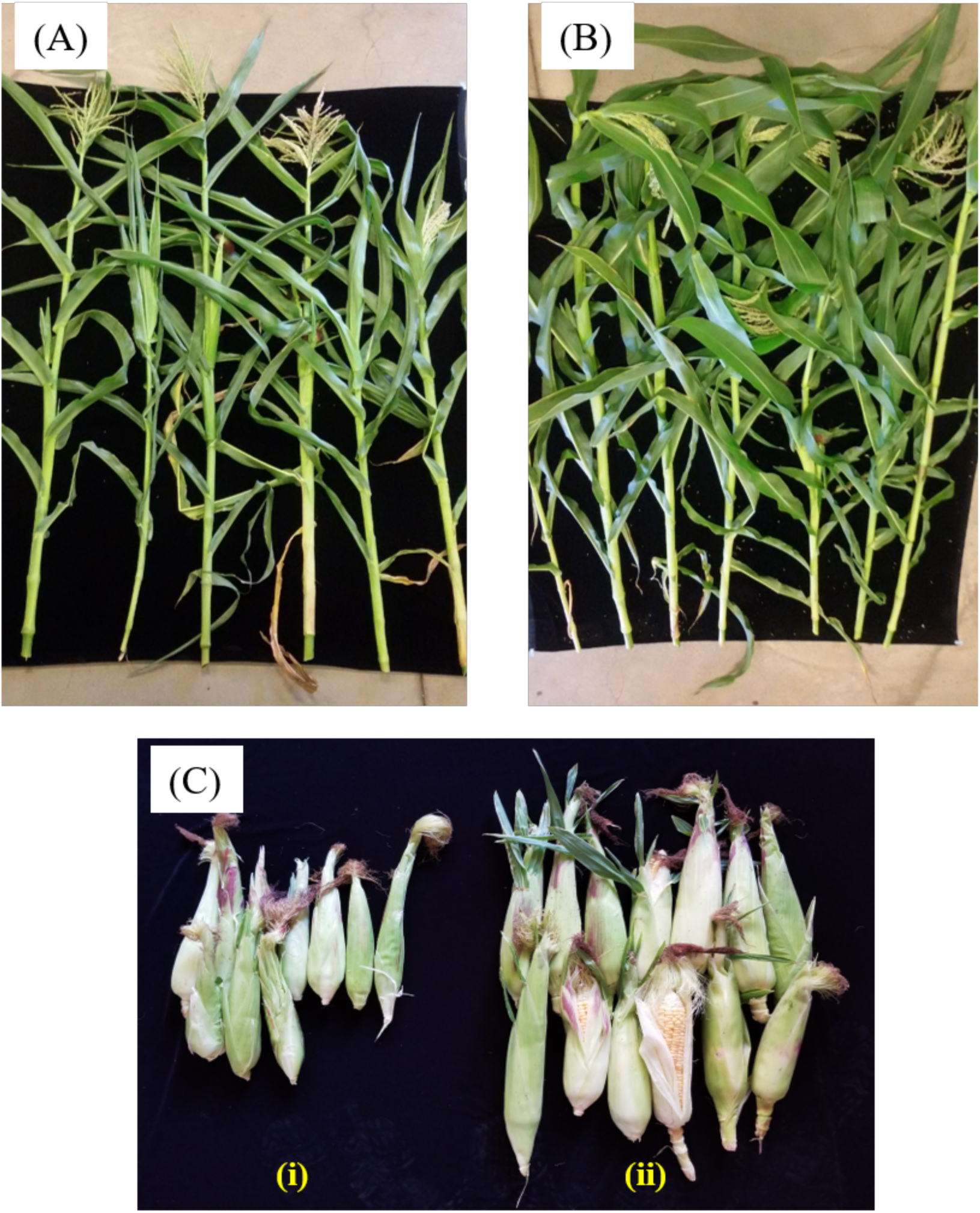
Impact of *D. cinnamea* 55 on the growth of corn (Outside garden trial). Figures (A) and (B) demonstrate the representative plants randomly chosen from the uninoculated control and *D. cinnamea* 55 treatment, respectively. (C-i and C-ii), harvest of control and treated sets, respectively.

**Table 2.** Effect of *D. cinnamea* 55 inoculation on growth of corn (Microplot trial)

| Parameters | Treatments |  |
| --- | --- | --- |
|  | Control | Treated |
| Shoot length (inches) | 33.71±10.01 | 47.19±6.21 |
| Dry shoot biomass (g) | 29.66±5.10 | 40.37±8.72 |
| Fresh wt. of 10 cobs (g) | 34.8±7.63 | 56.5±9.73 |
| Dry wt. of 10 cobs (g) | 17.9±6.54 | 37.3±5.77 |

Values are means of 24 plants ± SD for each of the treatments at the time of harvesting (after 120 days of sowing).

### Virulence tests with *C. elegans*

Assays with the nematode *C. elegans* are frequently used to analyze broad-host range microbial pathogenicity when *P. aeruginosa* PA14, which is an effective killer of nematodes, is used as a positive control. On NGM, *C. elegans* exposed to PA14 were motile, but avoided the bacteria, which remained unconsumed by the nematodes, which led to their death (Table 3) in contrast to their normal food source, *E. coli* OP50, which has no effect on worm viability. The test strain *D. cinnamea* 55 did not show any inhibitory effect on the motility and growth of worms (Table 3, Fig. 4).

**Fig 4.**
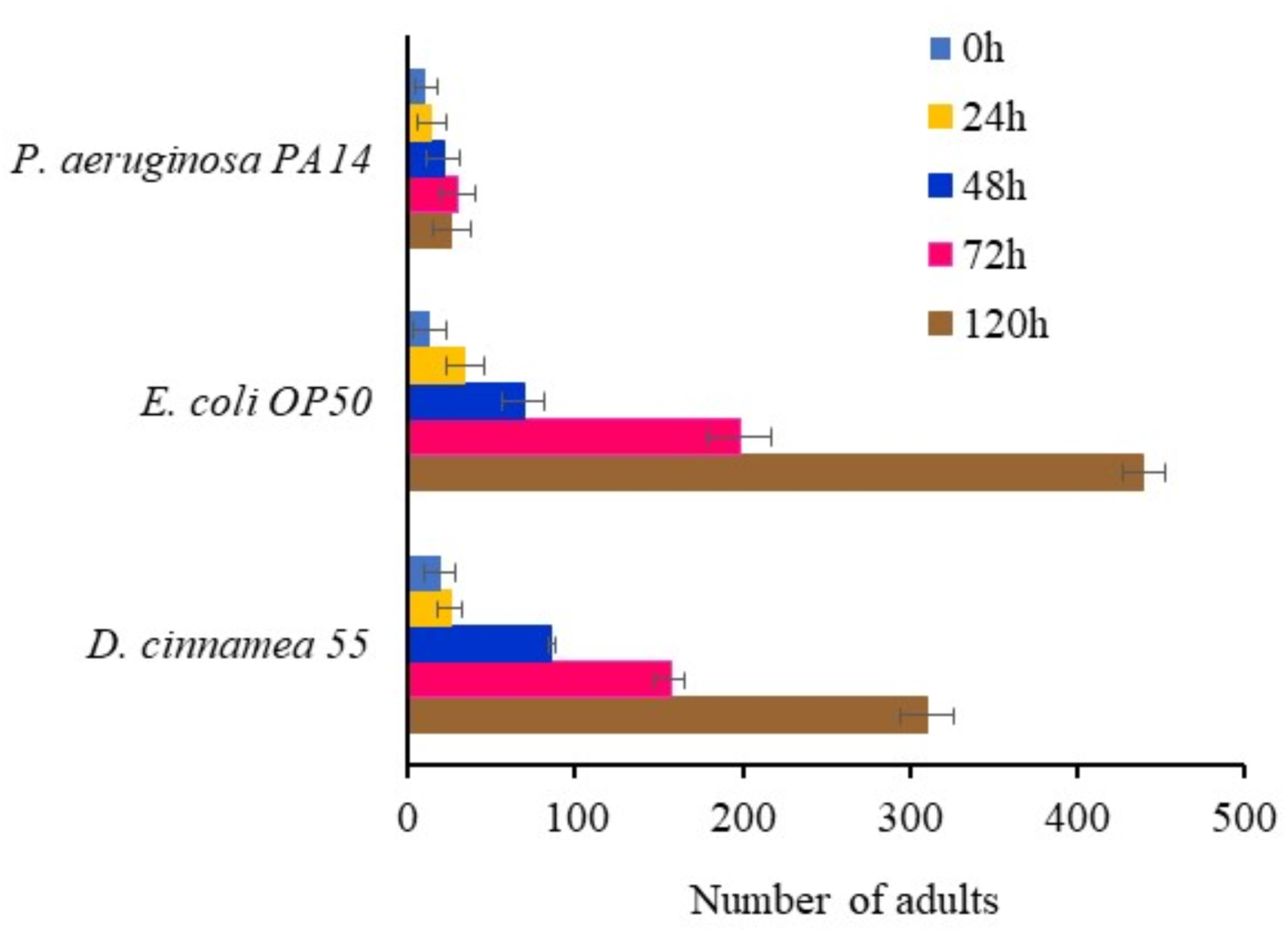
Response of *C. elegans* fed with *D. cinnamea* 55 under slow-killing conditions.

**Fig 5.**
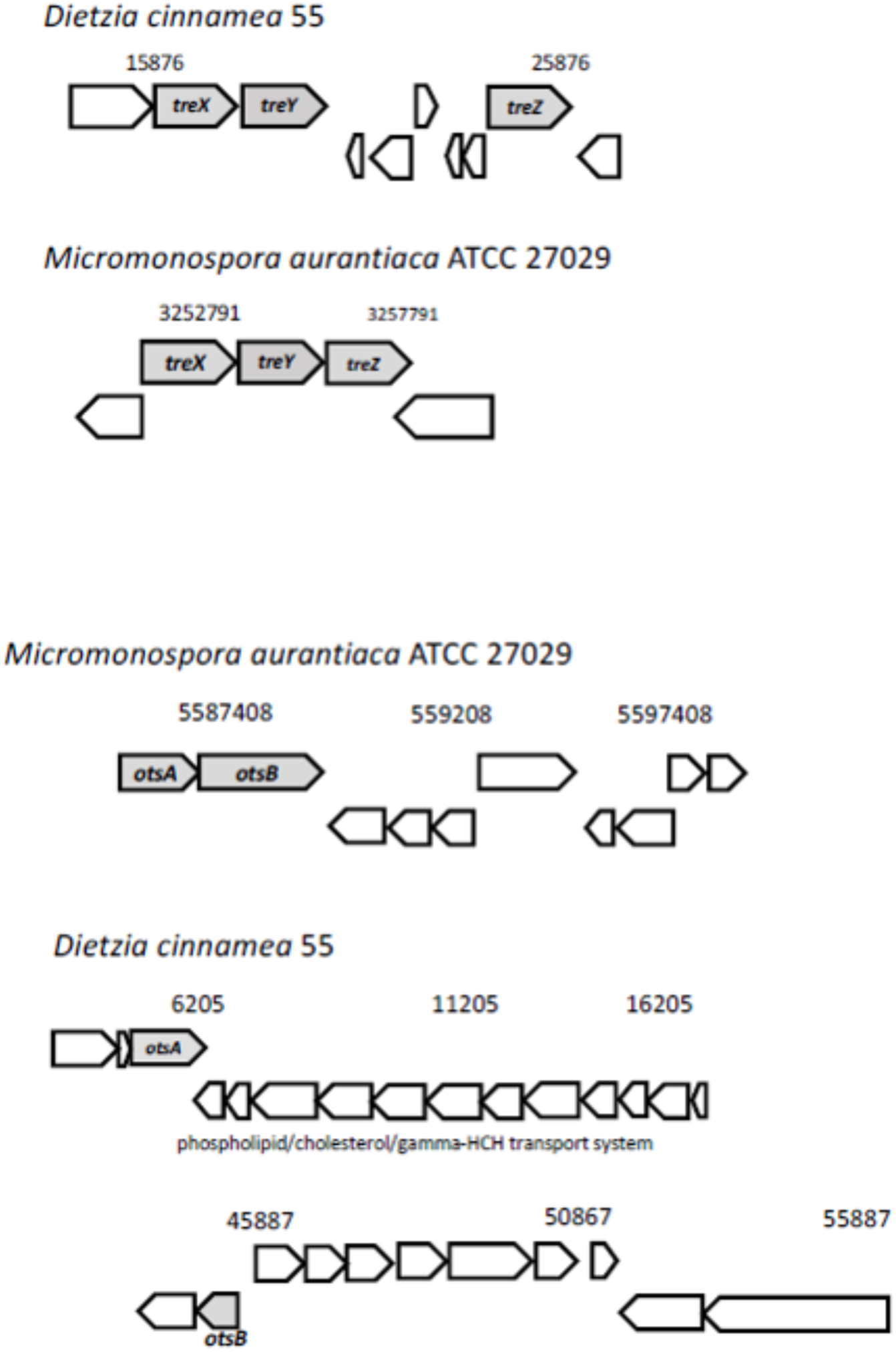
Possible mechanisms of stress tolerance via trehalose accumulation in *D. cinnamea* 55. The *treXYZ* genes in *D. cinnamea* 55 and *M. aurantiaca* ATCC 27029, MOTS pathway. The *otsA* and *otsB* are together in an operon for *M. aurantiaca* ATCC 27029 whereas in *D. cinnamea* 55, they are in two separate contigs, TPS pathway. The phospholipid/cholesterol/gamma-HCH transport gene system is shown for reference.

**Table 3.**
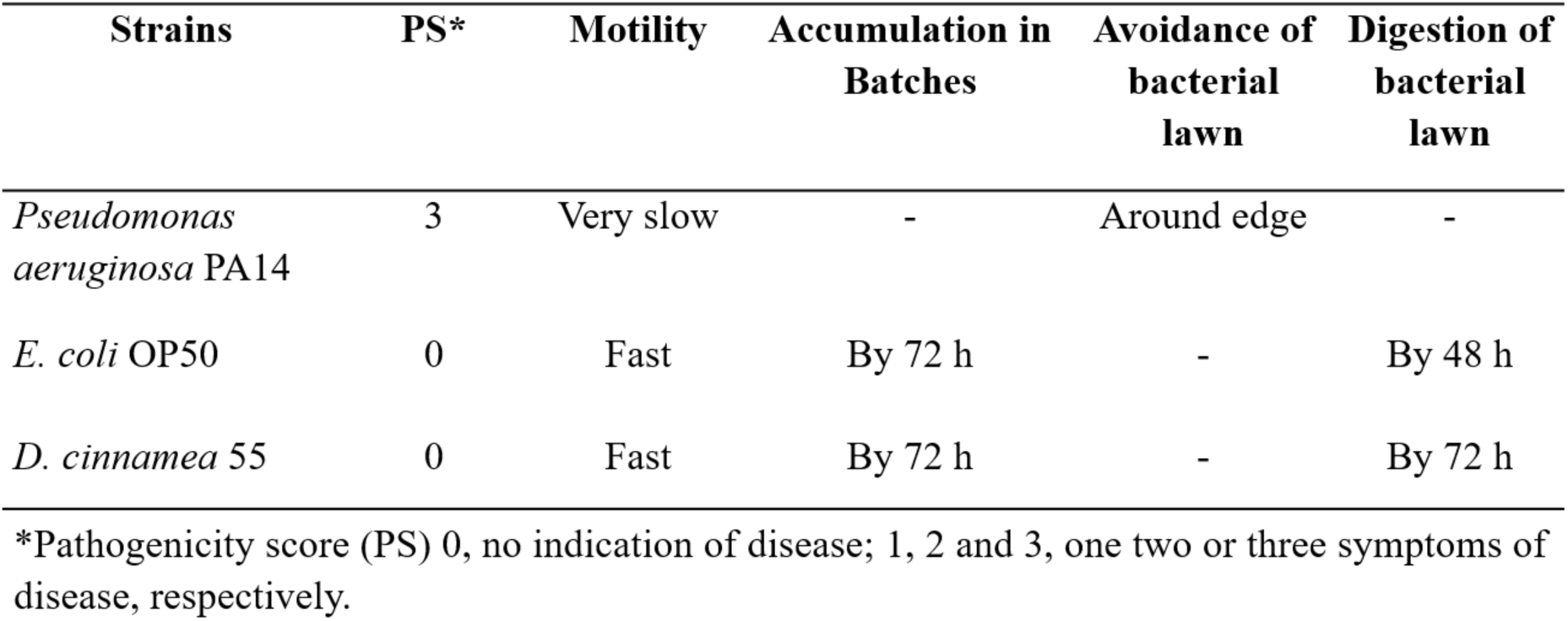
Pathogenicity score and behavioral response of C. *elegans* to the bacterial strains

**Table 4.**
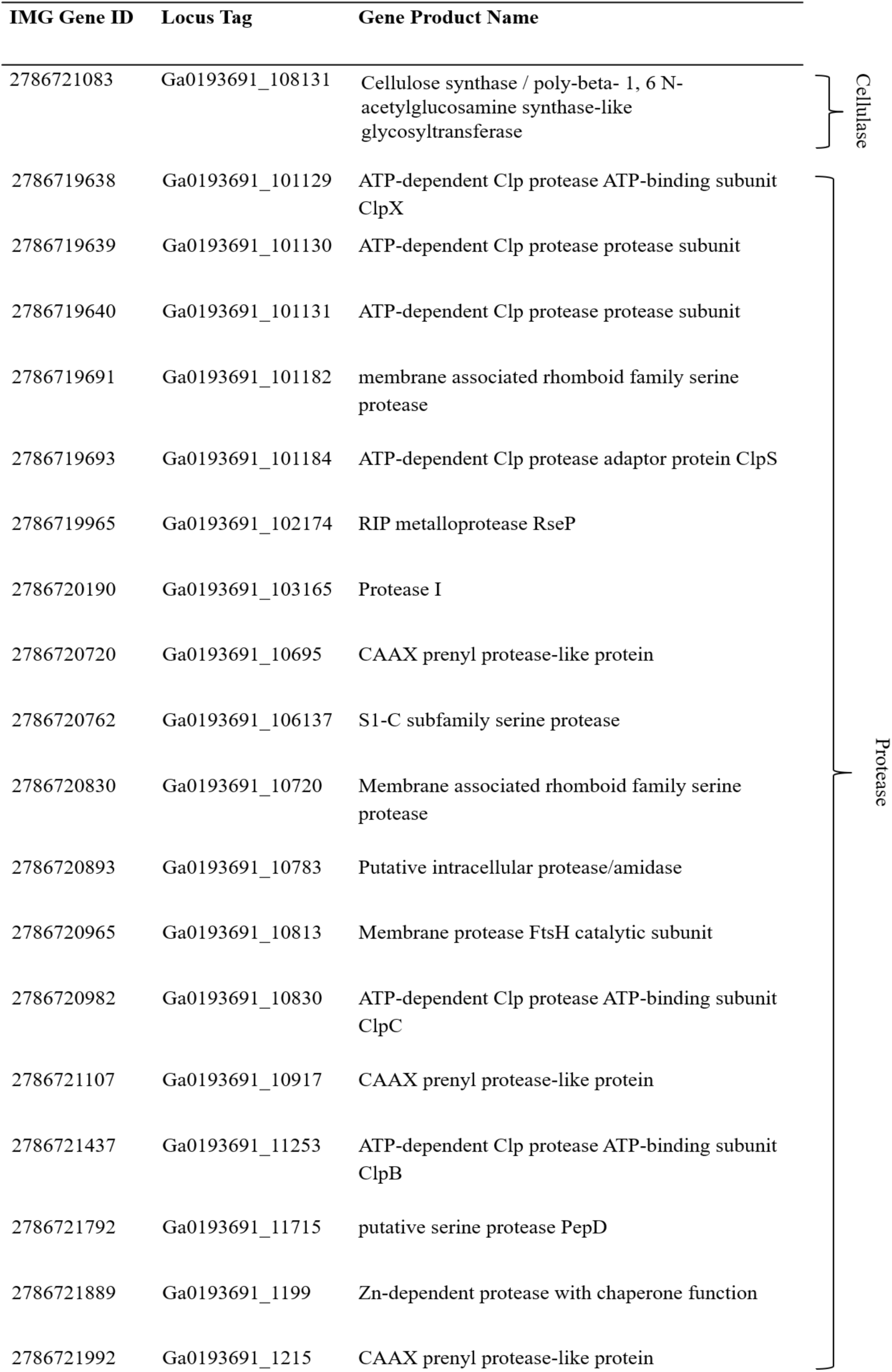
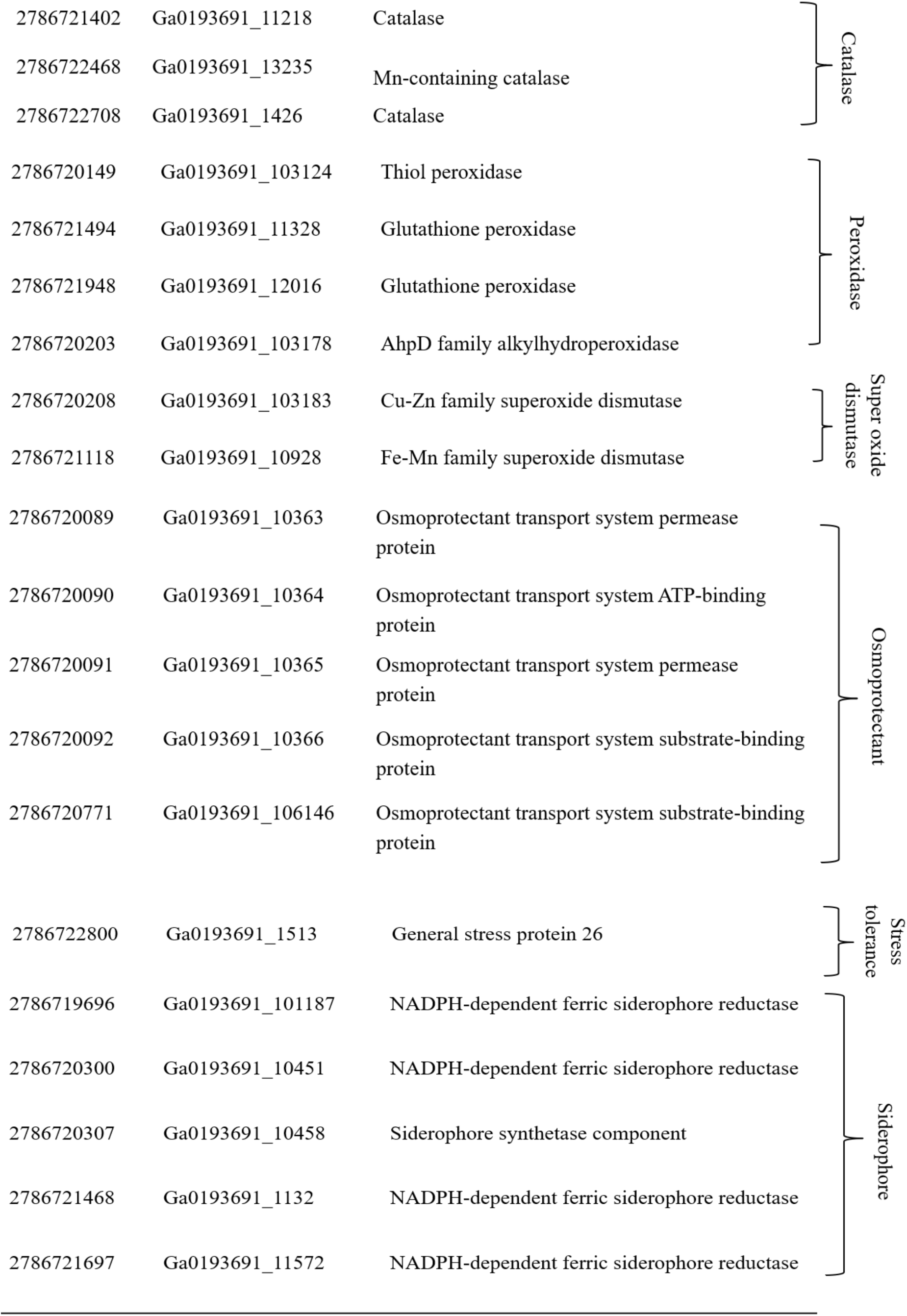
Plant beneficial function genes in *D. cinnamea* 55

The sequencing of the genome of strain 55 allowed us to look for gene sequences encoding potential virulence factors. The T3SS and T4SS, which are closely associated with virulence in Gram-negative plant pathogenic microbes, are not found in Gram-positive bacteria. Only one T3SS gene (IMG#2786720821; Ga0193691_10711) potentially encoding an inner membrane YOP/YscD-like protein and one T4SS gene (IMG2786722165; Ga0193691_124300), which if expressed would code for a TraM-binding TradD/TraG-like protein, were detected in the genome. On the other hand, genes for the T7SS are commonly found in Gram-positive microbes, and when expressed in pathogens, are responsible for virulence.

Queries for T7SS genes homologous to genes in *Mycobacterium tuberculosis* and *Staphylococcus aureus* yielded no hits in *Dietzia* strains. However, type VII secretion systems are found in bacteria that are not pathogens, for example, in model organisms such as *Bacillus subtilis*. Because they are less complex than those of *M. tuberculosis* or *S. aureus*, these systems are called Type VIIb secretion systems to distinguish them from the mycobacterial T7SS (Unnikrishnan et al., 2017). We queried the genomes of four plant-growth promoting bacteria namely, *B. subtilis* 30VD-1 and *B. simplex* 30N-5 for the presence of genes related to virulence. Both *B. subtilis* and *B. simplex* have a Type VIIb secretion system, which in *B. subtilis* 30VD-1 consists of 7 genes in contrast to the 20 different genes making up the *Mycobacterium* T7SS operon. The *yuk* genes encode an FtsK-SpoIIIE Atpase, WXG100 proteins, as well as other system-specific proteins.

*Dietzia* was found to be similar to *Rhodococcus equi* in several morphological traits (Koerner et al. 2009) and until the genus *Dietiza* was proposed by Rainey et al. (1995), many strains, both pathogenic and non-pathogenic, were named *Rhodococcus* (Yassin et al., 2006). Most genes related to virulence found in environmental non-pathogenic strains of Actinobacteria are conserved in the phyla, suggesting that pathogenicity is linked to certain acquired genes that trigger virulence (Letek et al 2010). Given this scenario, we looked for genes in the *Dietzia cinnamea* strain 55 genome that were relevant to the pathogenicity of *Rhodoccocus equi* and *Mycobacterium tuberculosis*, species related to *Dietzia*.

Virulence-associated protein A (VapA) is a protein encoded by a gene on a plasmid that is linked to pathogenicity in *R. equi*. Two polymorphic forms of the gene were found depending on PCR amplification from either virulent or environmental samples suggesting an important role of this protein in disease (Kalinowski et al. 2018). However, this gene was not detected in strain 55, implying a lack of virulence. Also, *R. equi* possesses six *mmpL* genes, encoding members of the “mycobacterial membrane protein large” family of transmembrane proteins, which are involved in complex lipid and surface-exposed polyketide secretion, cell wall biogenesis and virulence (Letek et al. 2010). Some non-pathogenic genera such as *Micromonospora* contain homologous genes, and *Dietzia cinnamea* 55 contains at least two of them: JGI id: 650236657, a superfamily putative drug exporter and the integral membrane protein *mmpl5* (JGI id: 650237246).

There are also four Fbp/antigen 85 homologs (REQ01990, 02000, 08890, 20840), involved in Mtb virulence as fibronectin-binding proteins and through their mycolyltransferase activity, are required for cord factor formation and integrity of the bacterial envelope (Letek et al. 2010). REQ01990 is not found in strain 55, and homolog 02000 (IS3 family transposase, partial), is a transposase but with only 50% identity to the *Mycobacterium* gene. The 08890 signal recognition particle-docking protein FtsY (*Mycobacterium tuberculosis*; gene ID 2786722813) has a higher identity, of 74% with a gene of the same name in strain 55, i.e. a signal recognition particle-docking protein FtsY (gene ID 2786719992).

## Discussion

Global climate change has accelerated the concurrence of a variety of abiotic and biotic stress factors with adverse effects on agricultural productivity (Batisti and Naylor, 2009; Jin et al. 2017; Meyers et al. 2017). A sustainable solution to this problem is through the use of root-associated microbes, which consist of numerous plant beneficial bacteria. The PGPB present an alternate technology for improving the stress tolerance capacity of plants (Khan et al., 2011; Vejan et al., 2016; Gamalero and Glick, 2015). We hypothesized that such bacteria were potential candidate for promoting plant growth and proceeded to isolate bacteria from under the canopies of indigenous plants living in the Negev desert (Kaplan et al., 2013).

We chose corn as a test plant because it is an important crop in American agriculture and faces a potential threat in production due to changing environmental conditions (Schlenkers and Roberts, 2009; Zhenong et al., 2017). A steadfast endeavor toward improving farming technology and management strategies as well as more basic research in agricultural microbes is needed for reducing the negative impact of climate change and its anticipated effects on corn yield. The present study demonstrates how the inoculation of abiotic stress-tolerant *D. cinnamea* 55 enhanced the overall plant health of corn in comparison to untreated controls. A more robust and developed root system was observed in the bacteria-treated plants. Stimulation of root growth and effective root area for enhanced water and nutrient uptake is a very important stress management tool because a healthy and proliferated root system helps the plant maintain optimal growth and development under stress conditions (Olanrewaju et al., 2019; Mendes et al., 2013). Both greenhouse and outside garden experiments show that *D. cinnamea* strain 55 had growth-promoting effects on corn as measured by shoot and root dry weight and also cob size and dry weight.

We found no evidence for virulence factors in this novel Gram-positive actinomycete based on both genome analysis and experiments on *C. elegans*. The lack of defined virulence signatures for *Dietzia cinnamea* 55 suggests that it is likely to be non-pathogenic. Indeed, *D. natronolimnaea* has been reported as a PGPB that protects wheat plants from salinity stress by regulating the expression of the plant’s stress-responsive genes (Bharti et al., 2016). Other *Dietzia* strains have been used for bioremediation, for biosurfactants, bioemulsifiers, and carotenoid pigment production, and also as therapeutic probiotics for ruminant animals with paratuberculosis (Click and Van Kampen, 2010; Click, 2011; Procópio et al., 2013; Gharibazhedi et al., 2013). Nevertheless, some *Dietzia* strains have been reported to be opportunistic pathogens (Hirvonen et al., 2012), which is the reason we focused on finding potential virulence factors in strain 55.

Surprisingly we could not detect obvious PGPB genes for plant growth promotion such as those found in other for phytohormone biosynthesis, other than the production of the polyamine spermidine, which interferes with ethylene biosynthesis (Maymon et al., 2015). Phosphate solubilization activity was not detected but siderophore production and also hydrolytic enzyme activity for potential entry into plant tissues were detected, suggesting that strain 55 could be an endophyte. The lack of the evidence in this study for the most typical PGP genes may result in part because the *D. cinnamea* 55 genome is unfinished. Another possibility is that protection from abiotic stress as conferred by strain 55 is sufficient for promoting enhanced growth at a crucial time in plant development, the germination and establishment stages. We are investigating this hypothesis further and will test *D. cinnamea* strain 55 on corn in a large-scale field study to verify its PGPB potential.

## Acknowledgments

This study was funded in part by grants from the Sol Leshin Program for Ben Gurion University-UCLA Academic Cooperation to D.K. and A.M.H. and from the Shanbrom Family Foundation to A.M.H. A CSP 1571 project from the DOE/JGI funded the genome sequencing of *Dietiza cinnamea* 55. Several UCLA undergraduates, C. Carmona, S. Pratap and S. Mohammadi, are acknowledged for their help with preliminary plant growth promotion assays.

We acknowledge the contributions of the late Dr. Yoav Bashan, a leader in the field of PGPB. He inspired us to pursue research in using microbes for soil restoration.

## Author contributions

AH, and NK were involved in planning and execution of the research, analysis and interpretation of the data, and writing the manuscript. PM-H, EH, and MM contributed intellectual advice and edited the manuscript. DK contributed strains and intellectual support.

---

**Fig S1.**
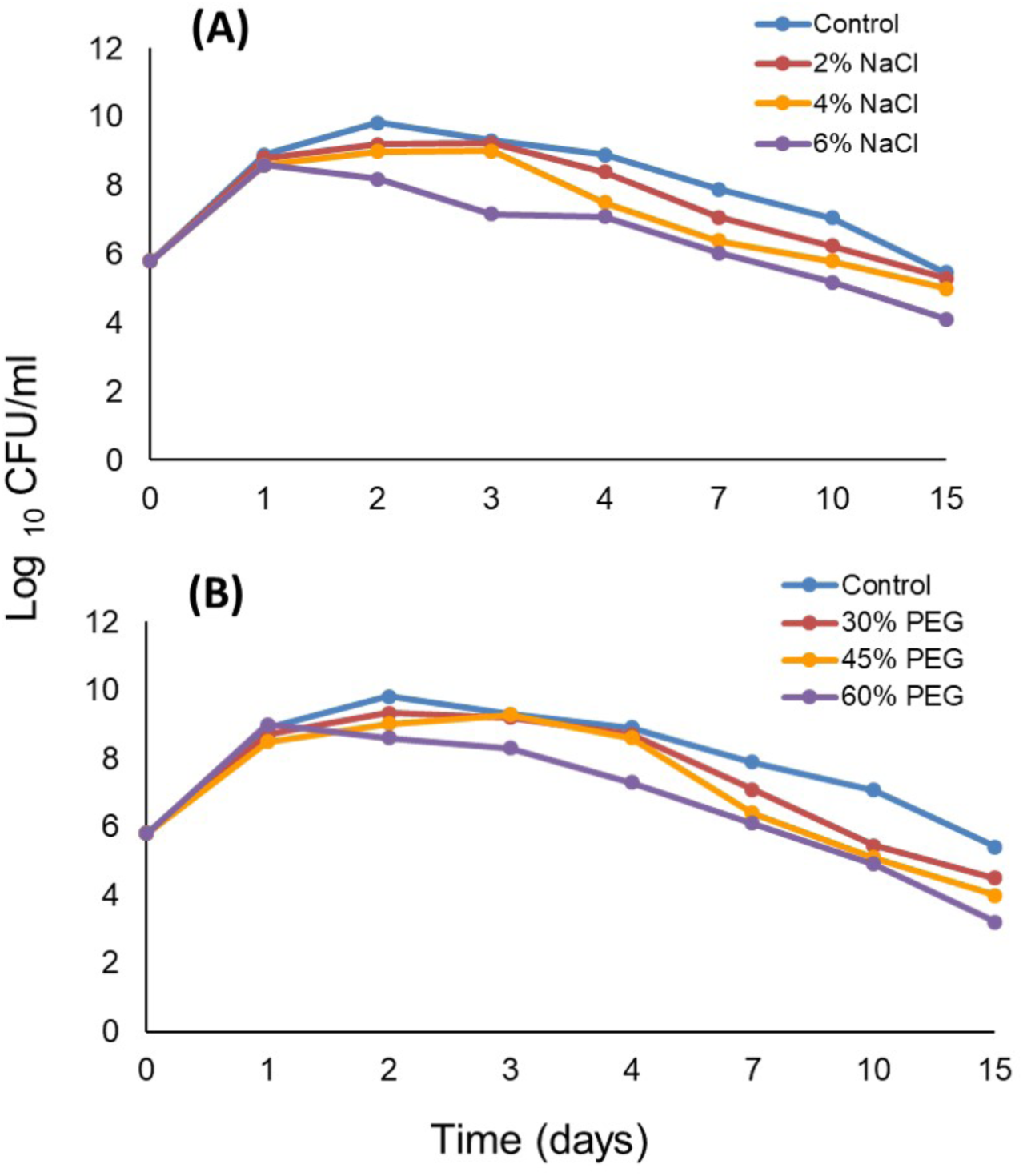
Salt (A) and PEG-induced drought (B) stress tolerance of *D. cinnamea* 55

**Fig S2.**
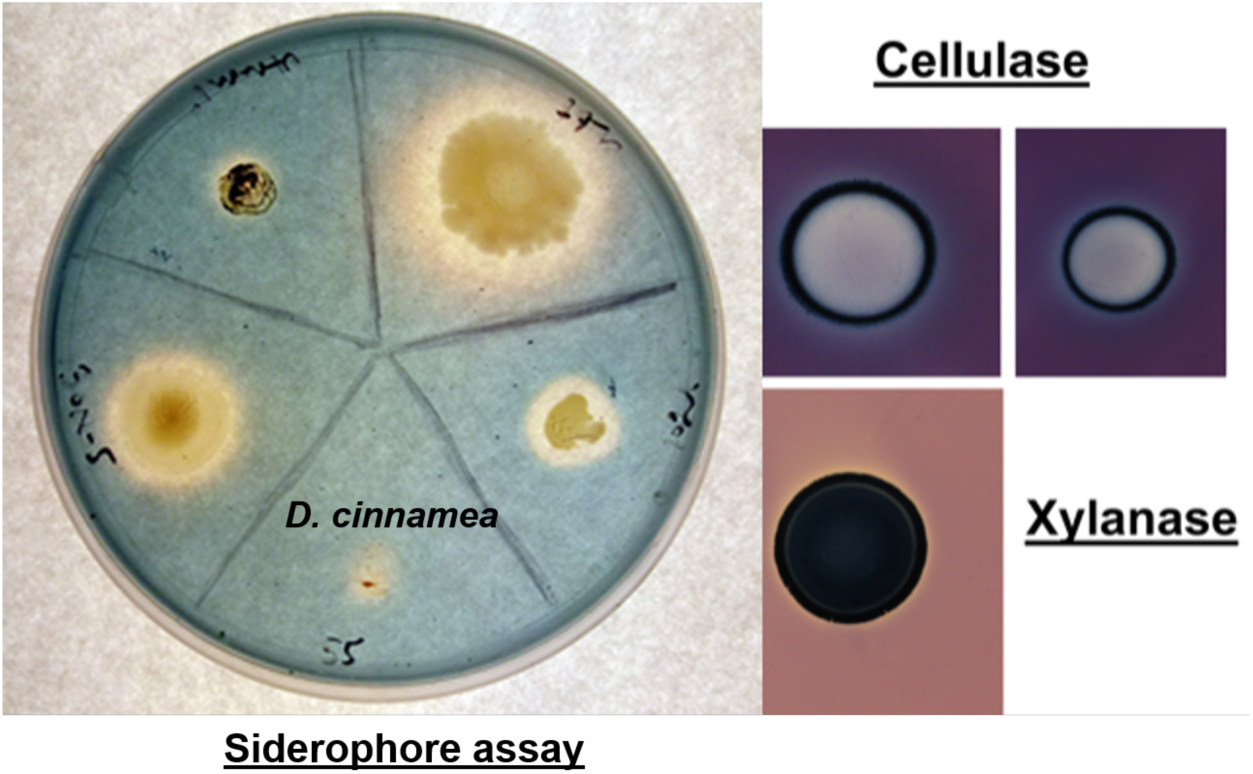
Photographs of siderophore production, and xylanase and cellulase activity of *D. cinnamea* 55

